# Differential alterations in peripheral tryptophan pathways in methamphetamine versus MDMA users are linked to their contrasting psychiatric symptoms

**DOI:** 10.1101/2025.08.25.672112

**Authors:** Francesco Bavato, Andrea Steuer, Anna M. Jacobsen, Amelie Zacher, Josua Zimmermann, David M. Cole, Antje Opitz, Markus R. Baumgartner, Ann-Kathrin Stock, Christian Beste, Boris B. Quednow

## Abstract

Methamphetamine (METH, “Crystal Meth”) and 3,4-methylenedioxymethamphetamine (MDMA, “Ecstasy”) are two types of substituted amphetamines that share structural-chemical similarities but exhibit contrasting acute and chronic effects including addictive liability. Tryptophan (TRY) pathways are involved in pleiotropic physiological functions at the interface of brain-body connections. Preclinical evidence suggests that amphetamines may modulate these pathways and, thus, indirectly influence brain functions via persistent alterations of peripheral metabolites. However, little is known about alterations of TRY-related metabolites in the blood and their clinical implications in chronic users of MDMA and METH. Hence, we characterized serum levels of TRY-related metabolites in a comparative cross-sectional study including n=36 chronic MDMA users, n=33 chronic METH users, and n=71 sex-matched, stimulant-naïve healthy controls (N_total_=140). An ultra– high performance liquid chromatography–mass spectrometry method was used to determine TRY metabolites. Combining metabolite levels, metabolic ratios, and network analysis we found robust evidence of divergent pathway alterations between METH and MDMA users. Chronic METH use was particularly associated with a depletion of serum TRY and serotonin levels, and a general activation of kynurenine pathways, while chronic MDMA use was linked to a selective activation of the OH-kynurenine metabolic branch. Metabolite changes were associated with the severity of psychopathology in the depression and psychosis domains across groups. Altogether, our findings demonstrate differential changes of serum TRY pathways in chronic MDMA and METH users. Persistent alterations of these pathways might contribute to the contrasting clinical profile of the substances and constitute a peripheral dimension of neurochemical plasticity with relevant implications for therapeutic targets.

## 1. Introduction

Methamphetamine (METH, “Crystal Meth”) and 3,4-methylenedioxymethamphetamine (MDMA, “Ecstasy”, “Molly”) are two types of substituted amphetamines that are widely used for recreational purposes in Europe and globally [1]. Despite their structural similarity and partially overlapping mechanisms of action, METH and MDMA differ markedly in their acute and long-term effects, as well as in their addictive liability. Both substances act as potent monoamine releasers, but they differ in their affinity for specific monoamine transporters: METH primarily promotes the release of dopamine (DA) and noradrenaline (NE), whereas MDMA predominantly releases serotonin (5-HT) and NE [2,3]. Chronic use of METH and MDMA induces neuroplastic adaptations in monoaminergic systems. METH has been associated with alterations in DA, NE, and 5-HT circuits, while MDMA has been more selectively linked to changes in serotonergic pathways, although with some species-dependent differences [4-7]. Putative neurotoxic effects and consequent structural brain alterations have been widely reported in both chronic METH users and MDMA users [8,9].

Clinically, chronic METH use is associated with affective symptoms (e.g., depressive symptoms and suicidal behaviour), psychotic-like experiences (e.g., paranoia, hallucinations, delirium, and delusion) increased impulsivity and violent behaviour, as well as high liability to addiction [3].

Consequences of chronic METH use also include dysfunctions in cognitive performance, decision-making, and empathy [10,11]. Notably, chronic METH use is also associated with relevant physical harm such as cardiovascular pathology (e.g., hypertension and myocardial pathology), reduced liver and kidney function, skin lesions (e.g., excoriations, ulcers, cellulitis), and poor oral hygiene (e.g. damaged and discoloured teeth, also called “meth mouth”) [3]. In contrast, chronic MDMA use is associated with a low risk of addiction and has limited relevance in clinical psychiatry, apart from transient depressive symptoms that often emerge a few days after use (a phenomenon referred to as “midweek blues”) [12]. However, whether chronic MDMA use can elevate the long-term risk for psychiatric disorders remains a matter of debate [13]. Like METH, chronic MDMA use has also been consistently associated with long-term cognitive deficits, although with higher domain-specificity manifesting as impairments in long-term memory and executive function, while socio-cognitive functions are usually preserved [14,15]. Despite decades of research, the mechanistic basis for the markedly different clinical outcomes of chronic METH and MDMA use remains insufficiently understood, particularly given their overlapping pharmacological profiles.

Peripheral 5-HT and tryptophan (TRY) pathways are involved in pleiotropic physiological functions at the interface of brain-body connections. Blood 5-HT levels have been shown to modulate sensory neuron activity and interoceptive signalling independently of 5-HT neurotransmission in the brain [16]. In this context, a reduction of blood 5-HT was causally linked to fatigue and cognitive impairments in the post-acute phase of viral infections. The 5-HT precursor TRY is an essential amino acid obtained exclusively through diet, critical for both multi-organ protein synthesis and brain-specific 5-HT metabolism through availability-limiting mechanisms [17]. Experimental TRY depletion in humans has been associated with depressive symptoms, memory impairment and increased aggression [18]. Circulating TRY catabolites produced in the liver and in immune cells (Kynurenine [KYN] metabolites) [19] or by gut microbiota (indole [IND] metabolites) [20] can cross the blood-brain barrier (BBB) and exert psychoactive functions by modulating glutamatergic transmission (via N-methyl-D-aspartate [NMDA] and α7 nicotinic acetylcholine receptors), microglial activation, and mRNA expression (via aryl hydrocarbon receptor signalling) [19]. In particular, the balance between kynurenic acid (KYNA) and hydroxy-kynurenine (OH-KYN), which are alternative products of KYN metabolism, has been found to shape neural-glial crosstalk, although only OH-KYN and KYN can directly cross the blood brain barrier, while KYNA is produced from KYN in the astrocytes [21].

Preclinical evidence indicates that substituted amphetamines may modulate TRY-related neurochemical pathways [22] and, thus, indirectly influence brain functions. However, in humans, little is known about how chronic MDMA and METH use affects blood levels of these metabolites, or how such alterations relate to clinical symptoms. Identifying whether METH and MDMA differentially affect TRY pathways could clarify the biological mechanisms underlying substance-related psychopathology and reveal potential targets for novel therapeutic interventions. Existing human studies have been limited by small sample sizes [23], comorbid medical conditions (e.g., HIV) [24], lacking investigation of KYN and IND metabolites [23-26], as well as limited documentation of substance use patterns and psychiatric symptomatology [23-26]. Moreover, previous studies only focused on METH with little to no investigation on MDMA, thereby precluding a clear understanding of the specific links between substance use, neurochemical alterations, and clinical outcomes.

Therefore, we aimed at characterizing serum levels of 5-HT, TRY, indole lactic acid (ILA) and indole propionic acid (IPA), KYN, KYNA, and OH-KYN in chronic METH and MDMA users. We hypothesized that chronic use of METH vs. MDMA would be associated with distinct alterations of the serum concentrations of these metabolites. We also conducted an exploratory network analysis to compare the organization of the metabolic pathway between MDMA and METH users in terms of edge weights, network centrality and eccentricity. Finally, we hypothesized that the alteration of peripheral metabolite levels would predict the distinct psychopathological profiles associated with the chronic use of each substance.

## 2. Methods

### 2.1 Participants

A total of N=203 participants were initially recruited between June 2019 and July 2021 by means of online media and flyer advertisements placed in institutions involved in substance use information and prevention, addiction clinics, and in public places. The final sample considered for this analysis included 36 chronic MDMA users and 36 age-, sex-, education-, verbal intelligence-, and nicotine use-matched stimulant-naïve healthy controls (HC) from Zurich, Switzerland, as well as 33 chronic METH users and 35 age- and sex-matched stimulant-naïve HC from Dresden, Germany. Exclusions from current analysis were based on lack of availability for blood samples (n=27), not fulfilling the requirements from substance use criteria according to toxicological data (n=31), relevant medical conditions (n=4), or pregnancy (n=1), according to previous publications [10].

All participants were aged 18 to 45 years. Substance users were eligible with a minimum of 25 lifetime occasions of MDMA or METH use and recent use within the last 12 months. Either MDMA or METH had to be the primary illicit substance for the respective group. General exclusion criteria included i) pregnancy or breastfeeding, ii) acute or previous severe somatic or neurological disorders, iii) severe DSM-IV Axis I psychiatric disorders such as autism and schizophrenia, iv) current intake of psychotropic medication (not applicable in the METH user group) and v) daily cannabis use (not applicable in the METH user group). Stimulant-naïve controls were excluded if they reported illicit substance use on more than 15 lifetime occasions (except cannabis) or intake of stimulants in the past 4 months (confirmed by hair analysis). All participants were asked to abstain from illicit substances for at least 48 hours (verified through urine substance screening) and alcohol for 24 hours before testing.

The study was approved by the Ethics Committees of the Canton Zurich (BASEC-Nr. 2018– 02125) and the TU Dresden (EK 69022018). All participants provided written informed consent and received compensation for their participation in accordance with the Declaration of Helsinki. Data on social cognition (whole sample) and brain imaging (MDMA sample only) from the same dataset have been published before [10,27,28].

### 2.2 Clinical and substance use assessment

All participants were screened for common DSM-IV Axis I disorders by trained examiners. Substance use disorders were additionally assessed with the Structured Clinical Interview for DSM-5 Axis I disorders. Subjective substance use history was further assessed with the standardized and structured Interview for Psychotropic Drug Consumption [29]. To objectively characterize substance use during the four months prior to testing, proximal hair samples of 4 cm were taken from the occiput. If a participants’ scalp hair length was not sufficient, body hair was sampled. Liquid chromatography-tandem mass spectrometry (LC-MS/MS) was used for toxicological hair analysis [30]. To additionally screen for recent illicit substance intake, multi-substance urine tests were conducted prior to the tasks. Self-report screening questionnaires were used to evaluate (i) symptoms of depression using the Center for Epidemiologic Studies Depression Scale (CESD) [31], (ii) attention-deficit/hyperactivity disorder (ADHD) with the ADHD Self-Rating Scale (ADHD-SR) [32], (iii) psychotic-like experiences with the Community Assessment of Psychic Experience (CAPE) [33]; and (iv) subjective trait impulsivity with the Barratt Impulsiveness Scale (BIS-11) [34]. To estimate the premorbid verbal intelligence (verbal IQ), the standardized German vocabulary test (Mehrfachwahl-Wortschatz-Intelligenztest) was applied [35].

### 2.3 Assessment of blood markers

Blood serum samples were drawn using silica and gel containing tubes (BD Vacutainer). All samples were collected in the afternoon (14-16:00) after a light meal. Samples were centrifuged for 15 min at 2000 rpm and subsequently stored at <−20 °C. Metabolite levels were analysed using an ultra–high performance liquid chromatography system (Thermo Fisher, San Jose, CA, USA), coupled to a 5500 linear ion trap quadrupole mass spectrometer (Sciex, Darmstadt, Germany), according to a well-established procedure from the same lab [36].

### 2.4 Statistical analysis

Statistical analyses were conducted in Python (version 3.9.6). Demographic and substance use data were analysed using Pearson’s chi-square tests and analyses of variance (ANOVA). All metabolite levels were winsorized in order to account for outliers that fell outside two interquartile ranges from the median. Metabolite levels were z-transformed per site using means and standard deviations (SD) of the respective HC sample to account for potential site-related differences in sex and diet [37,38]. Metabolic ratios were calculated to further quantify the general activation of the KYN pathway (KYN/TRY), of the serotonin pathway (5-HT/TRY), the alternative activation of the OH-KYN (OH-KYN/TRY) and KYNA (KYNA/TRY) branches and their relative balance (KYNA/OH-KYN) according to previous publications [39]. Group differences were assessed via linear mixed-effects model (LME) that predicted z-standardized metabolite levels or ratios with group and sex as fixed factors, random intercepts modelled for participants, and age and body mass index (BMI) as covariates. Sex, age and BMI were included considering their influence on TRY catabolites and 5-HT levels [37]. Pairwise comparisons for metabolites and ratios showing significant group differences were then assessed with Mann-Whitney U tests (to account for non-normal distribution and variance heterogeneity) and false discovery rate (FDR) correction. Effect size Cohen’s d was reported for group differences based on means and SDs. A sensitivity analysis additionally included urine toxicology results in the LME (METH users: amphetamine n=6, THC n=5, methamphetamine n=10, any psychoactive substance n=12; no urine positivity in MDMA users or controls) as a categorical predictor to control for potential acute substance effects. Additional separate LMEs included smoking (cigarettes/week) and alcohol use (grams/week) as continuous covariates. Dose-response relationships between cumulative substance use over the past 12 months and alterations in metabolite levels and metabolite ratios were investigated using LMEs corrected for sex, age and BMI in METH and MDMA users separately. Only metabolites and ratios showing significant group differences compared to HC were included in these analyses. Hair data were not used due to limited sample availability from the Dresden site.

To investigate further potential alterations in the organization of the metabolic pathways, explorative network analyses were performed using NetworkX [40]. First, correlation matrices across all z-standardized metabolites were calculated based on Spearman’s rank correlation coefficients for each group. Then the difference between correlation matrices for METH or MDMA users and their respective HC groups was calculated. These values were used as network edges, with each metabolite acting as a node. The following network parameters were computed: betweenness centrality, closeness centrality, eigenvector centrality, and eccentricity. This approach captures differential metabolic connections across groups by directly comparing all possible relationships between metabolites (MDMA vs. HC and METH vs. HC) and quantifying how they change across samples. Moreover, a paired t-test was performed to compare edges values across metabolites between separate networks (MDMA-HC vs. METH-HC).

Potential associations between metabolite levels, metabolite ratios, and psychopathology scores in the whole sample were investigated using LME models adjusted for age, sex, and BMI. FDR correction was used to control for multiple comparisons for each metabolite.

## 3. Results

### 3.1 Demographic and clinical characteristics

Demographic, clinical, and substance use characteristics are summarized in Table 1. Among demographic and clinical characteristics, significant group differences were identified in sex distribution, years of education, as well as ADHD-SR, CESD, BIS, and CAPE scores. Regarding alcohol, nicotine, and cannabis use, significant group differences were observed in self-reported consumption frequency, duration, and dose, with highest use in METH users. Substance use in HC was negligible and in line with inclusion and exclusion criteria.

**Table 1.**
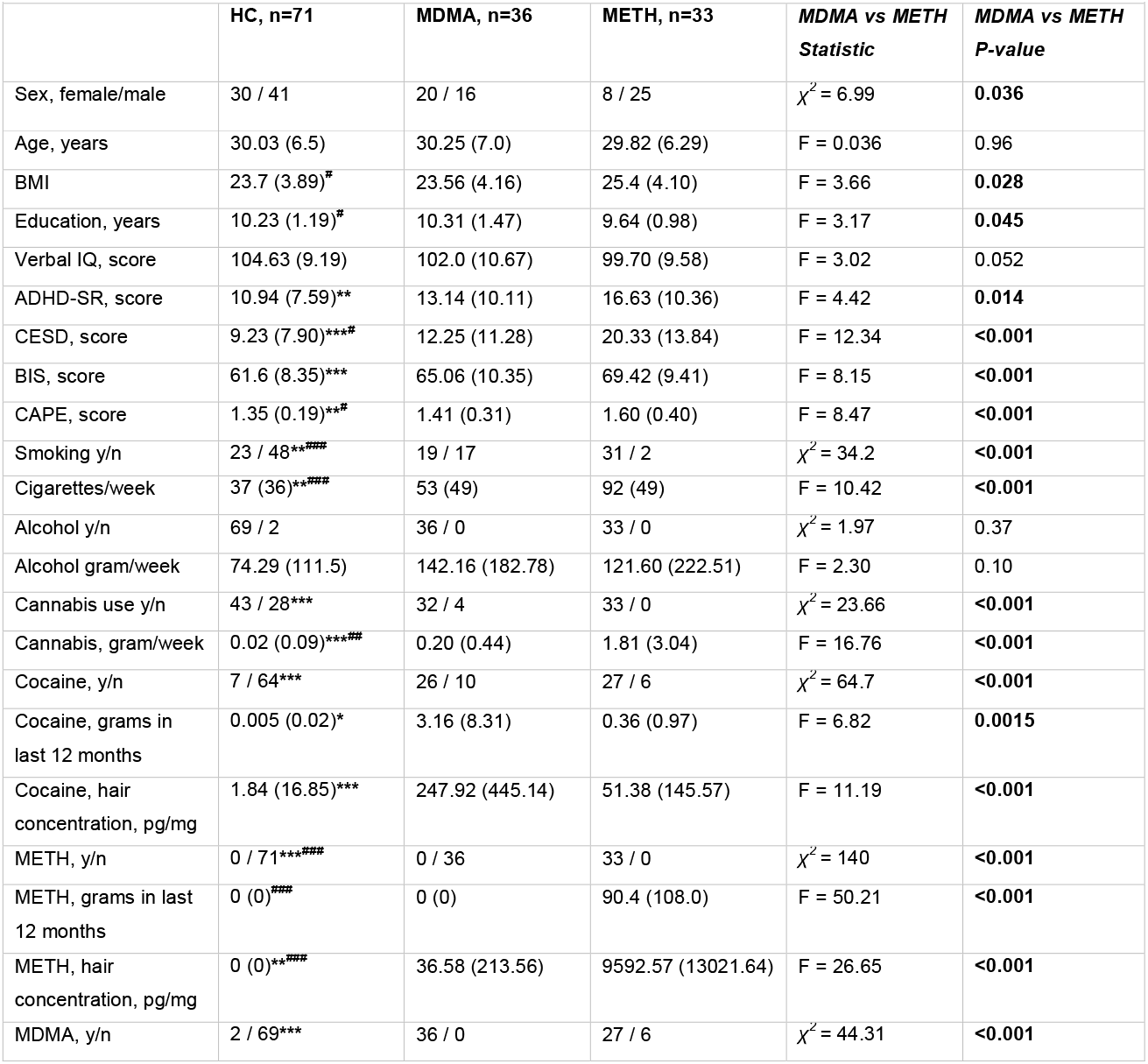
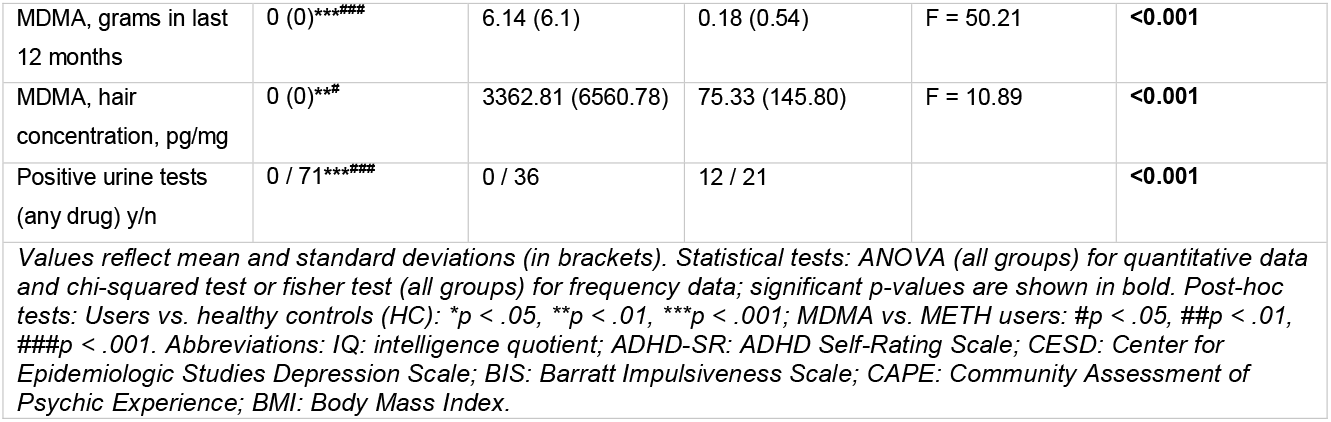
Demographic and clinical characteristics.

### 3.2 Group differences in serum metabolite levels and metabolite ratios

Serum metabolite levels and ratios across groups are reported in Figure 1. When adjusted for participant age, sex, and BMI using LMEs, group effects were found for 5-HT levels (p<0.001, d=0.45; pairwise comparisons FDR corrected: MDMA vs. METH: U=316, p=0.001, d=0.53; MDMA vs. HC: U=1128, p=0.32, d=0.22; METH vs. HC: U=1683, p=0.001, d=0.78) and TRY levels (p<0.001, d=1.03; pairwise comparisons FDR corrected: MDMA vs. METH: U=253, p<0.001, d=1.05; MDMA vs HC: U=1321, p=0.78, d=0.07; METH vs. HC: U=1749, p<0.001, d=1.05), demonstrating TRY and 5-HT reduction in METH users. LMEs also demonstrated KYN pathway activation in METH users (KYN/TRY, p=0.001, d=0.72; pairwise comparisons FDR corrected: MDMA vs. METH: U=708, p=0.26, d=0.38; MDMA vs. HC: U=1002, p=0.10, d=0.35; METH vs. HC:U=656, p<0.001, d=0.71), KYNA branch activation in METH users (KYNA/TRY, p=0.018, d=0.50; pairwise comparisons FDR corrected: MDMA vs. METH: U=760, p=0.07, d=0.58; MDMA vs. HC: U=1246, p=0.83, d=0.02; METH vs. HC: U=819, p=0.04, d=0.52), and OH-KYN branch activation in both METH and MDMA users (OH-KYN/TRY, p<0.001, d=0.85; pairwise comparison FDR corrected: MDMA vs. METH: U=617, p=0.78, d=0.04; MDMA vs HC: U=865, p=0.01, d=0.60; METH vs. HC: U=721, p=0.005, d=0.68).

**Figure 1.**
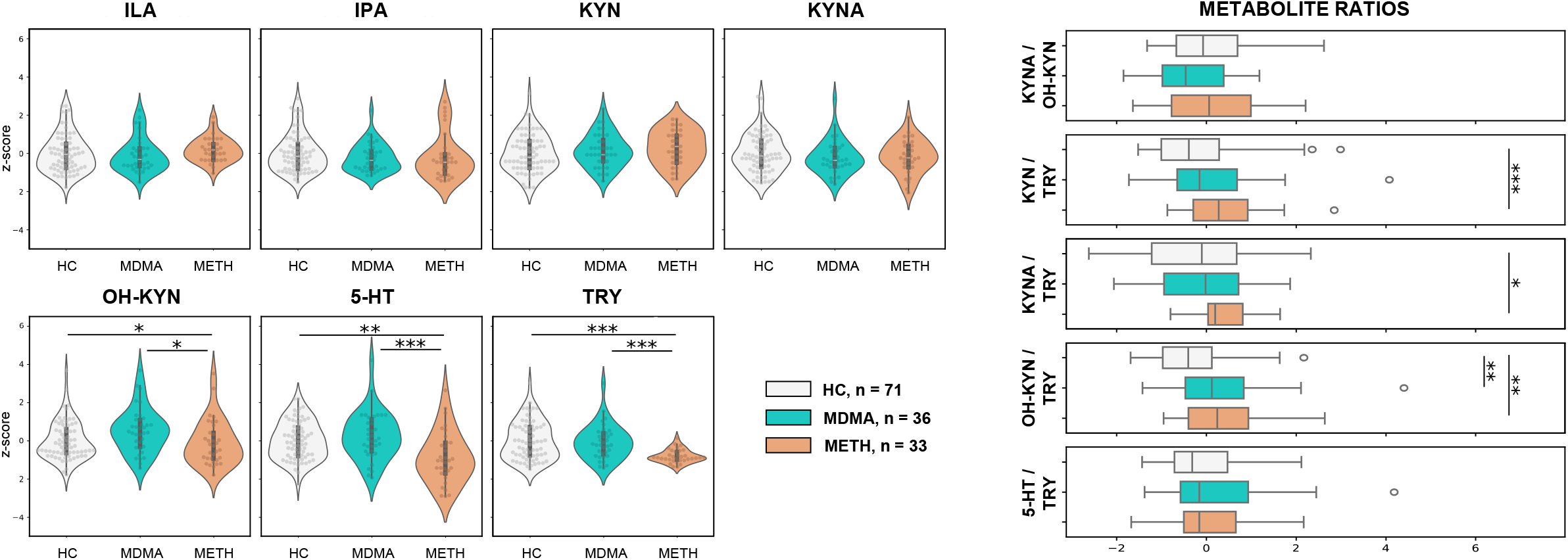
Left panel: violin plots with individual jitters showing metabolite levels (z-transformed) across samples. Right panel: boxplots showing ratios for metabolite levels across samples. Abbreviations: HC: Healthy Controls; ILA: indole lactic acid; IPA: indole propionic acid; KYN: Kynurenine; KYNA: kynurenic acid; OH-KYN: hydroxy-kynurenine; 5-HT: serotonin; TRY: tryptophan. *p < 0.05, **p < 0.01, ***p < 0.001.

### 3.3. Sensitivity analysis for serum metabolite levels

To clarify if the detected group differences (i.e., 5-HT, TRY, KYN/TRY, KYNA/TRY, and OH-KYN/TRY) are potentially dependent on acute or post-acute effects related to recent substance intake, we included urine positivity to any drug (urine positive cases in HC: n=0; MDMA users: n=0; METH users: n=12) as a factor in the LMEs. Urine positivity did not predict any of these values and ratios (all p>0.05). When considering urine positivity for METH specifically (METH users, n=10), a significant effect was found for 5-HT (p=0.006, d=1.75) but not for other values or ratios (all p>0.05). Adding urine positivity for METH in the LME, also weakened the group effect for 5-HT levels (p=0.225, d=0.24), suggesting this alteration to be rather acute/post-acute than chronic/persistent. In additional sensitivity analyses, neither alcohol use (grams per week over the last 12 months) nor nicotine (cigarettes per week) predicted 5-HT, TRY values or KYN/TRY, KYNA/TRY, or OH-KYN/TRY ratios in the LMEs (all p>0.05)

### 3.4 Exploratory network analysis

Exploratory network analyses of metabolic pathways are presented in Figure 2. Here, stronger alterations in network organization could be seen in the METH sample. In the edge matrix, mostly negligible to weak edge weight (correlation coefficient) contrasts (ρ=0.0-0.3) were found for the MDMA sample, while some moderate to strong contrasts (ρ=0.3-0.6) are observable in the METH sample (METH vs. HC). A global network estimation also visually confirmed significant differences in the metabolic pathways between MDMA and METH, with stronger alterations in edge weights in the METH sample (t=2.81, p=0.006). The most consistent differences between the two groups emerged in closeness centrality, which quantifies a node’s communication efficiency, and eccentricity, which quantifies distance to other nodes. Closeness centrality was higher across the network in the MDMA sample, while eccentricity was lower across the network in the MDMA sample than the METH sample.

**Figure 2.**
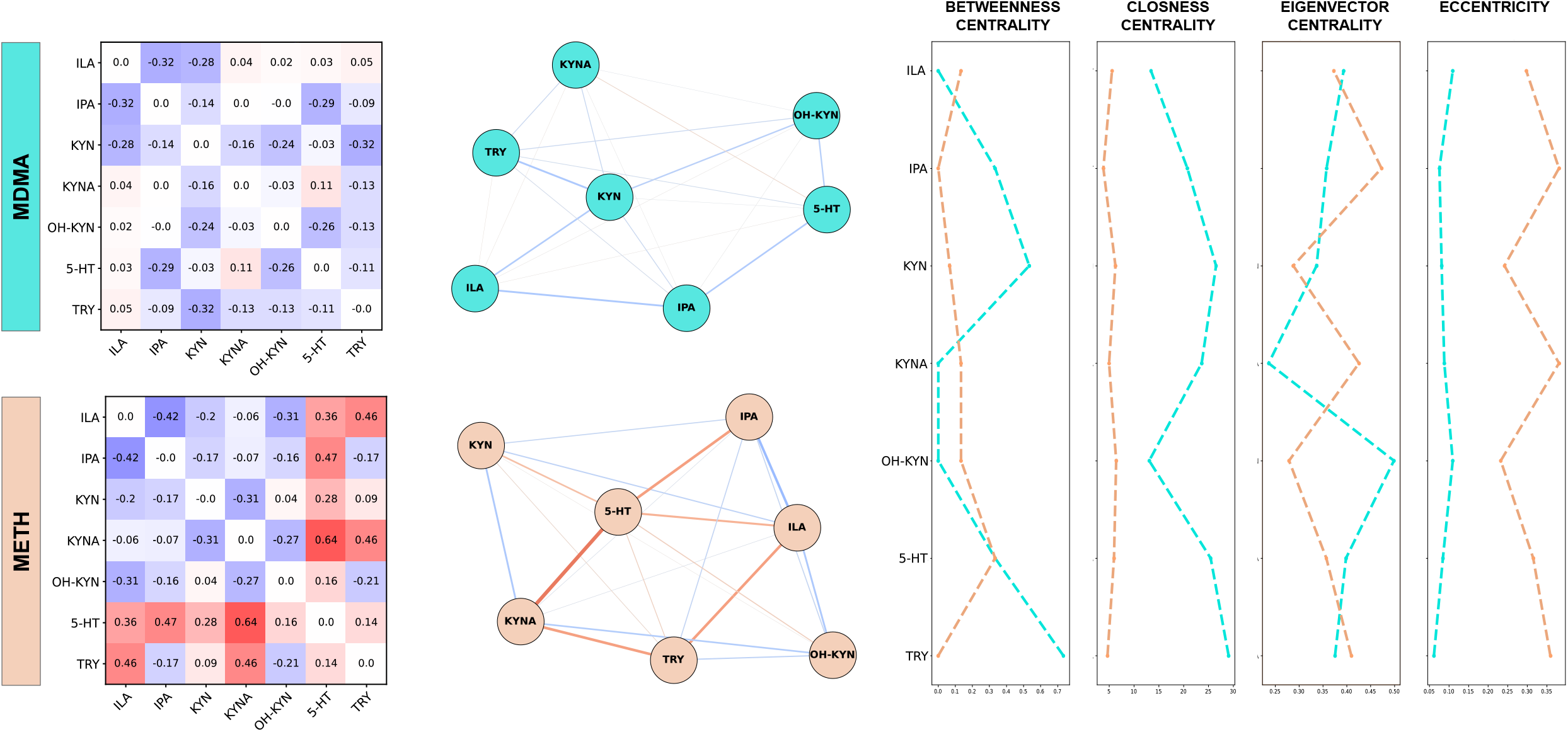
Left panel: correlation matrices across all z-standardized are calculated based on Spearman’s rank correlation as the difference between correlation coefficients between METH or MDMA users and healthy controls. Central panel: network visualization with metabolites as nodes and correlation coefficients as edges. Right panel: network parameters across nodes separated for each substance. Abbreviations: ILA: indole lactic acid; IPA: indole propionic acid; KYN: Kynurenine; KYNA: kynurenic acid; OH-KYN: hydroxy-kynurenine; 5-HT: serotonin; TRY: tryptophan.

### 3.5 Associations between blood markers and 12-month substance use variables

To address potential dose-response relationships, LMEs adjusted for age, sex, and BMI were used to detect associations between metabolites, ratios and self-reported METH or MDMA use in the last 12 months (see Figure 3). In the METH sample, a negative association was observed with TRY levels, while positive associations were found for the KYN/TRY ratio and OH-KYN/TRY ratio. A trend level negative association was observed between 5-HT levels and METH use. No significant associations were detected in the MDMA sample.

**Figure 3.**
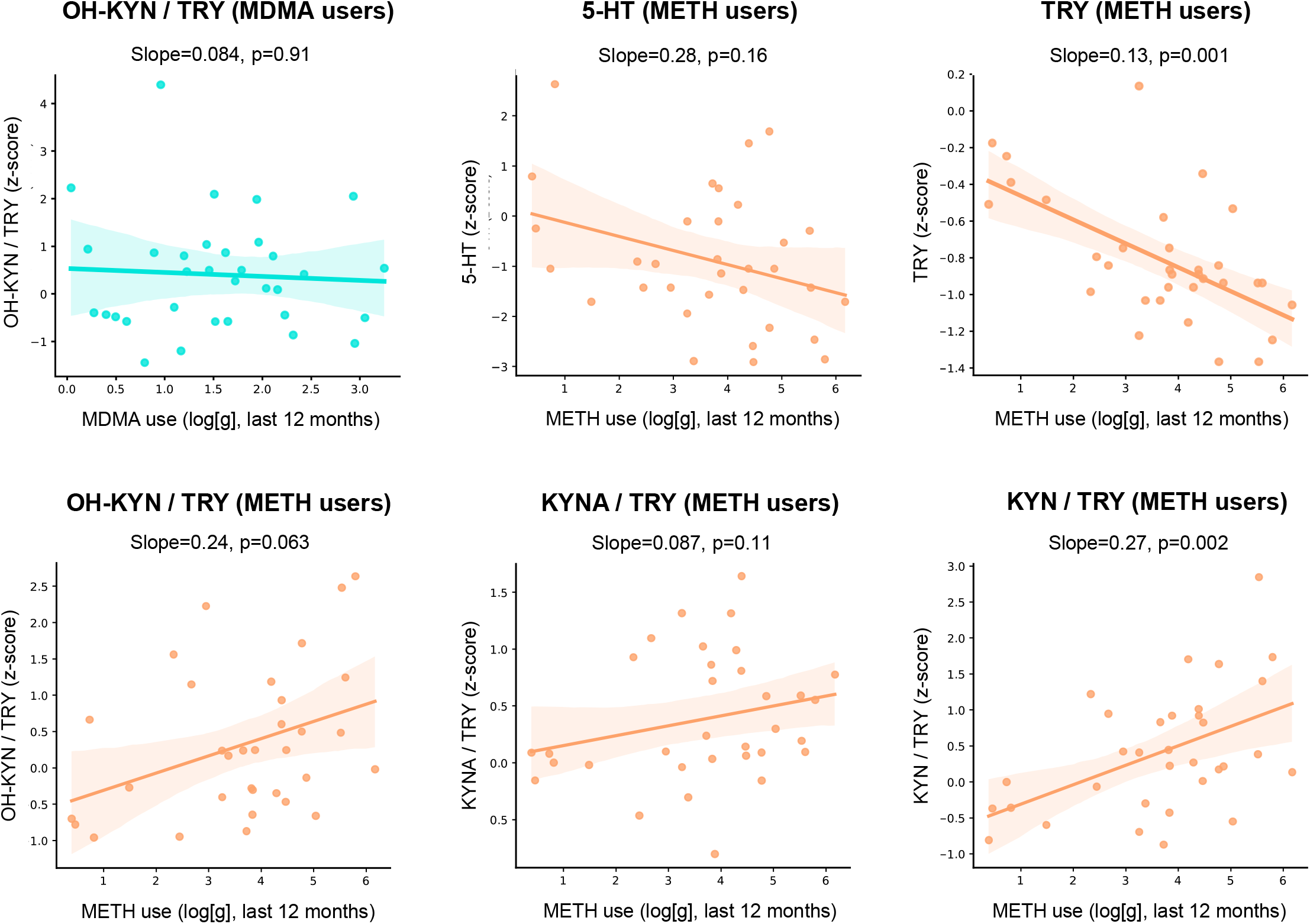
Scatterplots for dose-response relationships between metabolite levels (selected based on significant group effects) and self-reported substance use in the last 12 month. Mixed linear models adjusted for age, sex, and BMI were used. Abbreviations: KYN: Kynurenine; KYNA: kynurenic acid; OH-KYN: hydroxy-kynurenine; 5-HT: serotonin; TRY: tryptophan.

### 3.6 Associations between blood markers and psychopathology scores

To address potential associations of peripheral metabolites with severity in main self-reported psychopathology scores across the whole sample, LME corrected for age, sex, and BMI were performed (Figure 4). Significant associations (FDR corrected) were found between KYNA levels, CESD and CAPE scores. TRY levels were also associated with CESD and CAPE scores.

**Figure 4.**
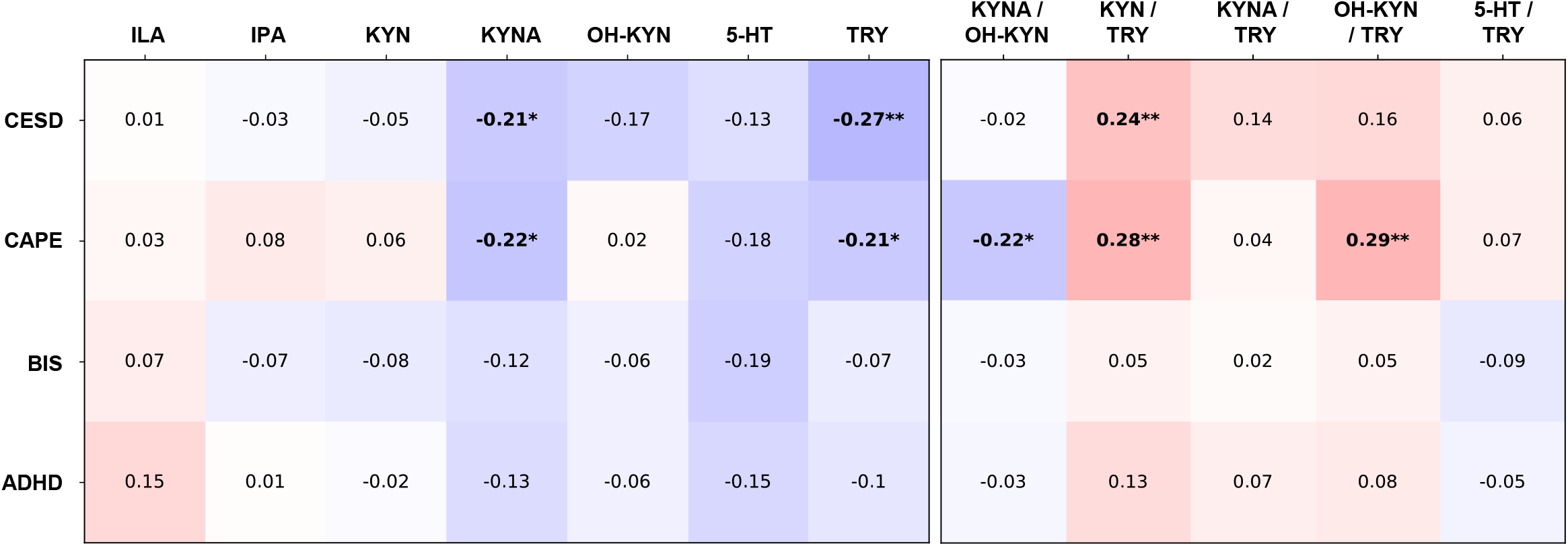
Heatmap reporting standardized beta values from mixed linear models adjusted for age, sex, and BMI. FDR correction was used to control for multiple comparisons for each metabolite. *p < 0.05, **p < 0.01.

Considering metabolite ratios, significant associations were found for KYN balance (KYNA / OH-KYN) with CAPE score, KYN activation (KYN / TRY) with CAPE and CESD scores, and OH-KYN branch activation and CAPE scores.

## 4. Discussion

The present study compared peripheral TRY pathways between chronic METH users, chronic MDMA users, and stimulant-naïve healthy controls. We provided robust cross-sectional evidence of differential alterations in metabolite concentrations and pathway organization across samples. In particular, METH users showed a reduction of TRY and 5-HT, a general activation of the KYN pathway, and moderate to strong metabolite network changes. On the other hand, MDMA was associated with only limited metabolite alterations. Metabolite levels and ratios were associated with severity scores in psychiatric symptoms in the total sample suggesting an involvement of peripheral TRY pathways in the differential clinical presentation of chronic use of METH vs. MDMA.

### 4.1 Chronic METH use is associated with serum TRY reduction

A depletion of blood TRY levels in METH users has been previously suggested, although studies to date employed only limited characterization of substance use variables (no toxicological data) and were highly heterogeneous in term of assays and matrices used [23-26,41]. Considering that TRY is exclusively acquired with diet, METH-related TRY reduction could either result from decreased input (i.e., dietary lack and malnutrition) or increased metabolism (i.e., conversion into the KYN pathway). Based on experimental evidence from preclinical studies (where substance-related malnutrition can be excluded), increased conversion to KYN is seen as the most probable cause [42]. This is also supported by the observed increase in KYN/TRY in our sample. Moreover, the association between METH use in the previous 12 months and both TRY and KYN/TRY in our study, support the suggested dose-response relationship of these effects. Considering the pleiotropic role of TRY, METH-related depletion might have a negative impact on protein synthesis across the body and also limit TRY-dependent 5-HT production in the brain. Notably, TRY pre-treatment in rats administered with METH was found to prevent METH-related behavioural changes [43]. On the other hand, experimental TRY depletion in humans was shown to elicit overlapping neurocognitive effects (i.e., impaired decision-making) with those observed in chronic METH users [44]. Thus, a peripheral TRY depletion may mediate the clinical phenotype of chronic METH use and represent a relevant target for treatment development. It should be noted that a previous study with experimental manipulation of dietary TRY failed to elicit pro-cognitive effects in MDMA users, however similar investigations in METH users (where blood TRY levels are more affected) are lacking [45].

### 4.2 Chronic METH use is associated with serum 5-HT reduction

The reduction of 5-HT levels in METH users is in line with previous preclinical studies. TRY metabolism into 5-HT is mediated by the rate-limiting tryptophan hydroxylase (TPH) enzyme. Both MDMA and METH have been widely shown to inhibit brain TPH activity and thus reduce brain 5-HT availability both acutely and long-term [46-48]. However, substance effects on systemic TPH have received far less attention. This distinction is particularly relevant as peripheral 5-HT does not cross the BBB. Accordingly, body and brain 5-HT are considered as parts of two distinct systems, with different biological implications. In our sample, reduced blood 5-HT might be a direct consequence of lower TRY availability, which is supported by unchanged 5-HT/TRY ratios across samples and normal 5-HT levels in MDMA users. Nonetheless, a modulation of THP activity beyond the brain is also possible. The observation of reduced serum 5-HT levels gains particular relevance in view of emerging evidence on the role of 5-HT in brain-body interactions [16]. Peripheral 5-HT has been recognized as a modulator of autonomic signalling via vagus nerve stimulation [49]. Blood 5-HT reduction has been causally linked to fatigue and neurocognitive functioning in post-viral syndromes, also via altered afferent nerve stimulation [16]. Moreover, body 5-HT levels modulate innate and adaptive immune responses and shape gut-immune interactions [50-52]. Some limited evidence has linked 5-HT-dependent immune-system alterations to mental health conditions such as affective disorders, OCD and autism [53-55]. Accordingly, alterations of body 5-HT pathways might be involved in chronic METH use. Importantly, METH effects on 5-HT in our study were associated with urine positivity suggesting a dependency on recent METH use rather than a long-term alteration. Thus, the interaction between METH and 5-HT metabolism may be reversible through abstinence.

### 4.3 Differential effects of MDMA and METH on the KYN pathway

While METH was associated with general activation of KYN metabolism (increase of KYN/TRY, KYNA/TRY, and OH-KYN/TRY ratios) but no alteration in the balance between the KYNA and OH-KYN branches, MDMA was associated with selective activation of the OH-KYN branch. Previous data on the effects of amphetamine derivates on KYN metabolites are very limited [41]. In rats, MDMA has been shown to induce an acute and subacute increase of KYN metabolism through indoleamine 2,3-dioxygenase activation (converting TRY to KYN) [56]. Modulatory activity of amphetamine derivates has also been suggested for the kynurenine aminotransferase (converting KYN to KYNA) while evidence for the kynurenine 3-monooxygenase (KMO, converting KYN to OH-KYN) is lacking [57]. Therefore, the activation of the OH-KYN branch in MDMA users is newly described and not clearly explained. Potential activation of the OH-KYN branch through KMO upregulation has been described for pro-inflammatory cytokines (i.e., interleukin-1β and IFN-γ), reactive oxygen species, and high O2 concentrations [58,59]. Notably, antidepressants (i.e., ketamine and selective serotonin reuptake inhibitors [SSRIs]) have been suggested to reduce (directly or indirectly) KMO activity [60,61]. Looking at the implications, OH-KYN crosses the BBB and elicits glutamatergic modulation (NMDA-R agonism) through its conversion into quinolinic acid. Neurotoxic effects of quinolinic acid have been widely described in preclinical studies and also suggested in humans [62,63]. Nonetheless, we did not observe consistent changes in the balance between the KYNA and OH-KYN branches across substances.

Overall, we demonstrated a different profile of KYN pathway alterations between METH and MDMA users, with relevant implications for their contrasting clinical presentation. On the contrary, no effects of MDMA and METH on IND metabolites were observed, thus suggesting a limited role of these gut-derived metabolites in the chronic effects of the substances.

Importantly, METH and MDMA also showed different organization of TRY pathways on a network level. While the sample size and the cross-sectional nature of our investigation prevent us from inferring a causal relationship on network directions, this explorative approach confirms the presence of differential effects of MDMA vs. METH on a global pathway level.

### 4.4 Associations between metabolites, ratios, and psychopathology

Our study provides novel evidence on the contribution of TRY pathways in shaping the clinical symptoms related to chronic MDMA and METH use. We particularly found that depressive symptoms (i.e., the CESD score) are negatively associated with TRY and KYNA levels, while being negatively associated with general activation of the KYN pathway. These findings are largely consistent with the literature on blood TRY levels in affective disorders [64]. Experimental depletion of blood TRY has been particularly associated with depressive symptoms through down-regulation of brain 5-HT production [18]. A potential pathway associating reduced blood 5-HT with affective symptoms through autonomic nerve signalling has also been proposed [16,49]. General activation of the KYN pathway through induction of the indoleamine 2,3-dioxygenase (i.e., by subclinical pro-inflammatory mediators and chronic stress) was also associated with depression [64]. Brain KYNA elicits NMDA-R antagonism and neuro-protective effects, thus KYNA reduction is also consistent with higher risk for depressive symptoms [21]. However, blood KYNA poorly crosses the BBB and its impact on brain KYNA is therefore questionable [65]. Nonetheless, blood KYNA alterations might still reflect an alteration in the KYN pathway which might elicit indirect effects on brain NMDA-R modulation.

Psychotic symptoms (i.e., CAPE score) were associated with decreased TRY, KYNA, and KYNA/OH-KYN ratio, and with increased KYN/TRY and OH-KYN/TRY ratios. These findings are contrary to the KYNA hypothesis of schizophrenia, which suggests that KYNA-dependent glutamatergic modulation shapes psychosis risk [66]. However, blood KYNA does not reflect brain KYNA levels and was found to have limited association with psychotic symptoms [67]. On the contrary, increased OH-KYN (which crosses the BBB) and OH-KYN pathway activation have been associated with psychotic symptoms in neuroleptic-naïve individuals with first-episode psychosis [68,69]. Here, non-linear modulatory effects on 5-HT and glutamatergic neurotransmission and the activation of neurotoxic pathways (via quinolinic acid activation) have been discussed as putative mechanisms.

### 4.5 Limitations

Our study bears some limitations. First, the cross-sectional nature of our investigation limits us in addressing the causal links between substance use and metabolite levels. In this context, the association between psychopathology scores and metabolite levels and ratios cannot disentangle effects of chronic substance use from pre-existing vulnerability. This is also limited by the intrinsic collinearity between psychiatric symptom scores and group membership, considering the higher severity in METH users. Future studies should include longitudinal data to better understand the dynamic course of this association and provide more robust evidence of dose-response relationships. Second, the lack of cerebrospinal fluid samples in the total sample and neuroimaging data in METH users do not allow us to assess potential associations between blood metabolites and brain functions. Third, chronic MDMA and METH use differ in use pattern and amount, thus affecting direct pharmacological comparisons. However, this reflects the difference in their real-world use and is therefore intrinsic in the naturalistic investigation of their contrasting clinical implications. Similarly, sex distribution was unbalanced in the user groups, which is also largely inevitable given that males are usually overrepresented in METH compared to MDMA user samples. Lastly, the moderate sample size might affect statistical power and significance detection at lower effect sizes.

Furthermore, future studies should include a characterization of inflammatory markers and enzyme activity to better clarify potential pathways involved. Nonetheless, we provide a robust demonstration of the contrasting effects of MDMA and METH on TRY pathways with relevant clinical implications.

### 4.6 Conclusions

The current investigation sheds light on the distinct alterations of peripheral TRY pathways in METH vs. MDMA users. We demonstrated substantial metabolite alterations in METH users (mostly driven by a reduction of TRY and 5-HT and an activation of the KYN pathway) and limited changes in MDMA users. We validated these findings using both case-control comparison of metabolite levels and an explorative network analysis approach. The metabolic changes showed relevant associations with psychopathology symptoms in the depression and psychosis domains, thus supporting the involvement of these pathways in the clinical presentation of chronic METH vs. MDMA use. Overall, our findings suggest that differences in clinical severity between METH and MDMA users may reflect not only central but also systemic metabolic alterations, pointing beyond a purely brain-focused view of psychoactive compound effects. Future studies should clarify the causal relationship between MDMA and METH use and changes in TRY-pathways and test experimental manipulations targeting these metabolites for treatment purposes.

## CRediT authorship contribution statement

**Francesco Bavato:** Conceptualization, **Formal Analysis**, Methodology, Writing – original draft, Visualization; **Andrea Steuer:** Methodology, Investigation, Writing – review & editing; **Anna Jacobsen: Formal Analysis, Visualization**, Writing – review & editing**; Amelie Zacher:** Formal analysis, Visualization, Writing – review & editing; **Josua Zimmermann:** Investigation, Data curation, Writing – review & editing; **David M. Cole:** Investigation, Writing – review & editing; **Antje Opitz:** Investigation; **Markus R. Baumgartner:** Investigation; **Ann-Kathrin Stock:** Investigation, Writing – review & editing; **Christian Beste:** Funding acquisition, Writing – review & editing; **Boris B. Quednow:** Funding acquisition, Conceptualization, Methodology, Writing – original draft.

## Acknowledgements

The authors would like to thank Samia Borra, Nicole Friedli, Mariana Carrillo Vázquez, Selina Caviezel, Marco Guglielmo, Xenia Häfeli, Mauro Mey, Anna-Katharina Schmid, and Babette Winter for assistance with recruitment and data acquisition.

